# Differential nuclear import determines lncRNA inheritance following mitosis

**DOI:** 10.1101/2022.10.06.511141

**Authors:** Michael D. Blower, Wei Wang, Judith Sharp

**Affiliations:** Department of Biochemistry, Boston University School of Medicine. 72. E. Concord St. Boston MA. 02118

## Abstract

Mitosis results in a dramatic reorganization of chromatin structure in order to promote chromatin compaction and segregation to daughter cells. Consequently, mitotic entry is accompanied by transcriptional silencing and removal of most chromatin-bound RNAs from chromosomes. As cells exit mitosis, chromatin rapidly decondenses and transcription restarts as waves of differential gene expression. However, little is known about the fate of chromatin-bound RNAs following cell division. Here we explored whether nuclear RNA from the previous cell cycle is present in G1 cells following mitosis. We found that half of all nuclear RNAs are inherited in a transcription-independent manner following mitosis. Interestingly, the snRNA U2 is efficiently inherited by G1 cells while lncRNAs NEAT1 and MALAT1 show no inheritance following mitosis. We found that the nuclear protein SAF-A, which is hypothesized to tether RNA to DNA, did not play a prominent role in nuclear RNA inheritance, indicating the mechanism for RNA inheritance may not involve RNA chaperones that have chromatin binding activity. Instead, we observe that the timing of RNA inheritance indicates a select group of nuclear RNAs are reimported into the nucleus after the nuclear envelope has reassembled. Taken together, our work demonstrates that there is a fraction of nuclear RNA from the previous cell cycle that is reimported following mitosis and suggest that mitosis may serve as a time to reset the interaction of lncRNAs with chromatin.

## Introduction

During cell division, the structure of the nuclear genome is dramatically altered to facilitate chromosome segregation to daughter cells. As cells enter mitosis, interphase chromatin loops and intrachromosomal interactions are dissolved and replaced by a series of nested loops created by the condensin complex [1, 2]. Concomitant with this mitotic chromatin remodeling, nuclear RNA transcription by all three polymerases is dramatically reduced [3]. In addition, nascent transcripts [4] and many components of the transcription machinery are removed from mitotic chromosomes.

During interphase, hundreds to thousands of RNAs (both coding and noncoding) are retained in the nucleus where they influence chromatin structure and gene expression [5]. At prophase, the vast majority of nuclear RNA is removed from chromatin [6], including notable nuclear RNAs such as XIST and repetitive lncRNAs [7–9]. Our group recently found that Aurora-B dependent phosphorylation of the nuclear RNA/DNA-binding protein SAF-A is responsible for the removal of the vast majority of nuclear RNAs from chromatin during mitosis [6]. Consequently, the RNA content of mitotic chromosomes consists chiefly of nucleolar RNA in the perichromosomal layer [10–12], and lncRNAs arising from centromeres and pericentromeres[10, 11].

At the completion of mitosis, recently segregated chromosomes decondense and are rapidly incorporated into a newly reformed nuclear envelope. Proper nuclear formation requires establishing crosslinks between chromatin [12, 13] and correlates with dissociation of the perichromosomal layer from chromosomes [14]. Following nuclear envelope reformation, nuclear proteins are reimported into the nucleus and transcription restarts. However, it is not known if nuclear RNA from the previous interphase is also reimported into G1 nuclei. Previous work has suggested that XIST RNA and repetitive RNA is not inherited in G1 cells [7, 8]; while pre-rRNA from the previous cell cycle is reincorporated into the nucleolus in G1, where it resumes processing and maturation [15]. However, it is not known how different types of nuclear RNA are inherited in G1 or if different pathways regulate the process of nuclear RNA inheritance. In this work we examine the behaviors of nuclear RNA following mitosis and find that nuclear import discriminates between different classes of nuclear RNA in early G1.

## Results and Discussion

### A fraction of nuclear RNA is inherited following cell division

To determine whether nuclear RNAs are inherited by daughter cells following mitosis we used EU [16] to follow the behavior of RNA in dividing cells. We synchronized DLD-1 cells in G2 by treatment with RO3306 [17] for 24 hours, and then labeled all cellular RNA by incubating G2-arrested cells with EU for 3 hours. Under these labeling conditions, the vast majority of RNA staining in G2-arrested cells is present in the nucleus with prominent staining of the nucleoli. Cells were then released from the G2 block and EU was removed from the culture media. We assessed nuclear RNA staining 1.5 hours after release from RO3306 and identified cells that had recently divided by staining for Aurora-B accumulation in the midbody, a marker for cells that have recently undergone cytokinesis (Fig. 1A). We then quantified total nuclear RNA signal in G1 cells compared to G2 cells that have not divided. Under this quantification scheme, inheritance of 100% of nuclear RNAs by daughter cells in G1 would result in 50% of the G2 signal. In our experiments we found that G1 cells had approximately 25% of the G2 nuclear signal, indicating that about half of nuclear RNA is inherited by G1 cells following division (Fig. 1B). The signal in telophase/G1 cells was significantly greater than that observed in cells lacking EU labeling (Fig. 1B). We observed some bright globular concentrations of EU RNA in G1 nuclei that were colocalized with nucleolar markers (Figure S1A), consistent with the observation that pre-rRNA present in the nucleolus at mitotic onset is inherited by G1 cells to complete processing [15].

**Figure 1.**
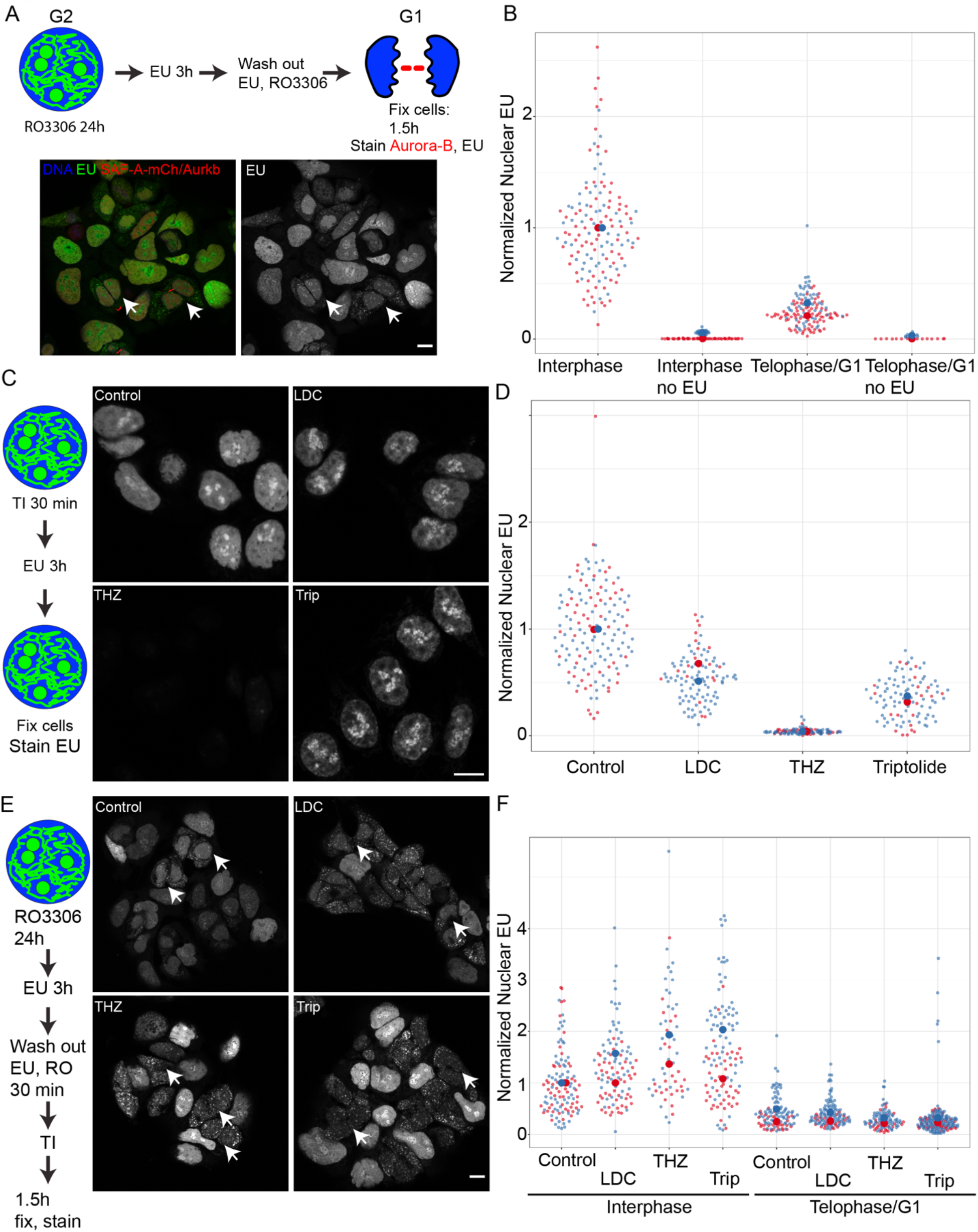
A fraction of nuclear RNA is inherited following mitosis. A. Cells were arrested in G2 using RO3306, followed by RNA labeling with 5-Ethnyl-Uridine (EU) for 3 hours. Cells were released from G2 and passed through mitosis. G1 cells were identified using Aurora-B staining of the midbody (arrow). Nuclear EU signal was quantified in G2 and G1 cells. B. Superplot [30] depicting the normalized nuclear EU fluorescence in two biological replicates. C. Transcription inhibitors were added to asynchronous cells for 30 minutes, followed by 3 hours of EU labeling. EU was detected following the labeling period. D. Plot of nuclear EU fluorescence from two biological replicates of each transcription inhibitor. E. Cells were synchronized and labeled with EU in G2 as described in A. Cells were released from G2 and transcription inhibitors were added in mitosis to block G1 transcription. Transciption Inhibitor (TI) F. Nuclear EU was quantified in G1 cells compared to G2 cells for each transcription inhibitor. There were no significant differences in G1 nuclear EU fluorescence. Scale bars are 5 μm for all images.

To determine if nuclear RNA inheritance is affected by the restart of genomic transcription in G1 cells, we examined the effects of a panel of RNAPII inhibitors. We examined the effects of a CDK9 inhibitor that blocks transcription elongation (LDC000067) [18], a CDK7 inhibitor causing transcriptional pause and release (THZ1) [19], and an XBP inhibitor that blocks initiation (Triptolide) [20]. We first validated the effects of each transcriptional inhibitor by testing EU incorporation and detection in asynchronous cell populations. Cells were incubated with each transcription inhibitor for 30 minutes, followed by 3 hours of labeling with EU (Fig. 1C). We then detected EU and measured total nuclear EU signal compared to untreated controls. We found that LDC and Triptolide significantly reduced global levels of EU signal while maintaining strong EU incorporation into nucleoli, consistent with specificity for RNAPII (Fig. 1CD). In contrast, THZ1 completely blocked all EU incorporation throughout the nucleus, suggesting that THZ1 can inhibit transcription by multiple RNA polymerases. To determine if transcription affected the reimport of EU labeled RNA in the nucleus after nuclear envelope reformation, we added transcription inhibitors to cells after release from G2 (Fig. 1E). Interestingly we found that transcription inhibition during mitosis and G1 did not affect the appearance of EU labeled RNA in G1 nuclei (Fig. 1F). We conclude that a fraction of nuclear RNA is inherited by G1 nuclei following mitosis and is independent of transcriptional restart in G1.

### Distinct classes of nuclear RNA exhibit differential inheritance patterns in G1

To gain insight into what types of RNA make up the nuclear EU RNA in G1 cells we examined several recent studies that identified transcripts physically associated with chromatin [21, 22]. We collected the lists of chromatin-enriched transcripts from two different studies using different methodologies and a total of three different cell lines. Interestingly, we found very little overlap in the chromatin-bound RNAs identified in these studies (Fig. 2A). Differences in identification of chromatin-bound RNAs were primarily attributable to purification methodology. In total, we identified 42 transcripts that bound to chromatin in both studies and all three cell lines. These chromatin enriched transcripts contained a large proportion of sn-, sno- and sca-RNA, as well as many well-validated nuclear lncRNAs such as NEAT1 [23], MALAT1 [24], and hnRNPU-as1 (Fig 2B). This informatic analysis suggested that different classes of RNAs are present on chromatin and may exhibit differential behavior during mitosis and G1.

**Figure 2.**
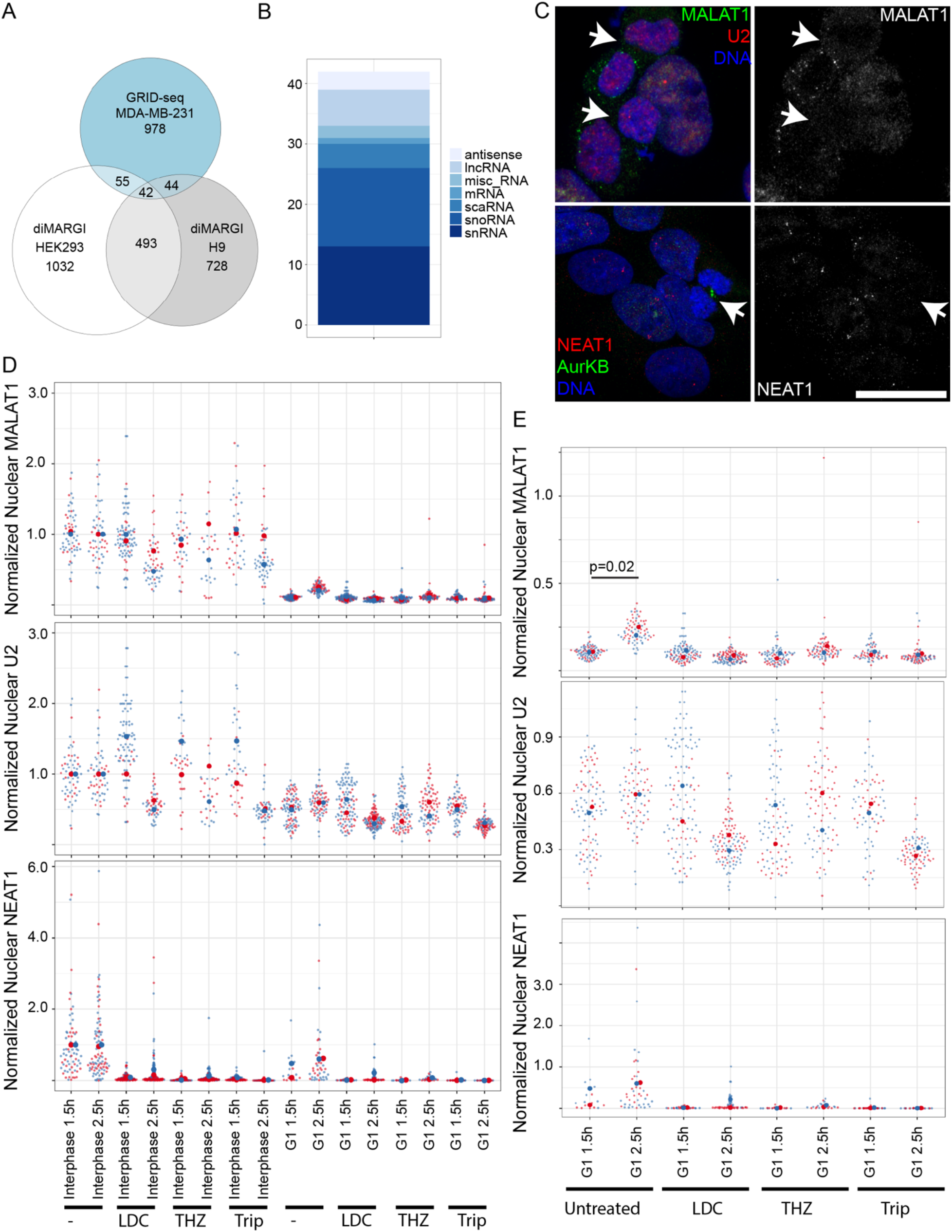
Different classes of nuclear RNA are enriched on chromatin and exhibit differential inheritance in G1. A. Chromosomal RNAs identified in two different genome-wide studies from three different human cell lines were compared to find conserved chromosomal RNAs. Venn diagram depicts overlap of RNAs identified in these studies. B. Biological classes of each of the 42 conserved chromosomal RNAs from A. C. Cells were synchronized in G2 and released into G1 as described in Figure 1. Three different nuclear RNAs from two different classes U2, MALAT1, and Neat1 were detected by RNA FISH in G1 cells. D-E. Cells were synchronized in G2 as described in Figure 1, then released into mitosis where they were incubated with different transcription inhibitors for either 1.5 or 2.5 hours. Nuclear RNAs were detected by RNA FISH and the amount of nuclear RNA FISH fluorescence was quantified in G2 and G1 cells. Plots depict the amount of nuclear RNA FISH signal for each RNA in G2 and G1 cells after treatment with various transcription inhibitors. All statistically significant changes are indicated on the plot. All G1 conditions were compared pairwise using a Wilcoxon-rank sum test. Scale bars are 10 μm.

To examine the inheritance of individual RNAs representing the snRNA and lncRNA classes we performed RNA FISH in early G1 cells for U2, MALAT1, and NEAT1 (Figure 2B, S1B). We synchronized cells in G2 by treatment with RO3306, then examined recently divided cells 1.5 or 2.5 hours after release from G2. Interestingly, we found that nuclear U2 RNA levels accumulated to the same level as G2-arrested cells very early in G1 (Figure 2CD). In contrast, MALAT1 and NEAT1 transcripts exhibited very low nuclear levels in early G1, which increased only after an additional hour in G1 (Figure 2DE). Additionally, we could identify clear cytoplasmic signal for MALAT1 and NEAT1 in early G1 cells while it was virtually impossible to detect cytoplasmic U2 signal (Figure 2C). Collectively, these results suggest that U2 RNA is inherited following mitosis while the lncRNAs NEAT1 and MALAT1 are not.

To determine if the low levels of nuclear MALAT1 and NEAT1 observed in G1 cells resulted from a low level of nuclear reimport or from G1 transcription we examined the effects of mitotic transcriptional inhibition on RNA inheritance. We synchronized cells in G2, released them into mitosis, then inhibited transcription using LDC, THZ1 or Triptolide. We allowed cells to proceed into G1 for 1.5 or 2.5 hours followed by detection of individual transcripts by RNA FISH. We detected rapid nuclear accumulation of U2 RNA in early G1 independent of transcription (Figure 2DE). In contrast, transcription inhibition during mitosis/G1 completely blocked nuclear accumulation of MALAT1 and NEAT1 RNAs (Figure 2E). Taken together, these results demonstrate that the U2 RNA is inherited by daughter cells following mitosis while the lncRNAs NEAT1 and MALAT1 require G1 transcription to accumulate in the nucleus. This result suggests that control mechanisms are in place to differentially sort whether lncRNAs undergo nuclear RNA inheritance following mitosis.

### SAF-A plays a minor role in nuclear RNA inheritance following mitosis

We recently reported that the nuclear factor SAF-A, which is hypothesized to play a role in tethering RNA to chromatin in interphase [25], is phosphorylated in the DNA-binding domain by Aurora-B during mitosis to trigger global RNA removal from mitotic chromatin [6]. We found that mutation of the Aurora-B phosphorylation sites (S14, S26) to alanine caused retention of RNA on mitotic chromosomes and chromosome segregation defects. To test the role of SAF-A in nuclear RNA inheritance following mitosis, we examined the effects of SAF-A-AA and SAF-A depletion on global RNA inheritance. We found that SAF-A depletion using SAF-A-AID resulted in no changes to global nuclear RNA inheritance following mitosis despite complete loss of SAF-A from cells (Figure 3A, S2AB) [6]. In contrast, we found a small but reproducible increase in nuclear RNA inheritance in cells expressing SAF-A-AA (Figure 3B, S2C). Collectively, these results demonstrate that SAF-A is not required for the normal pattern of RNA inheritance, but that ectopic retention of SAF-A on mitotic chromatin leads to modestly increased nuclear RNA inheritance.

**Figure 3.**
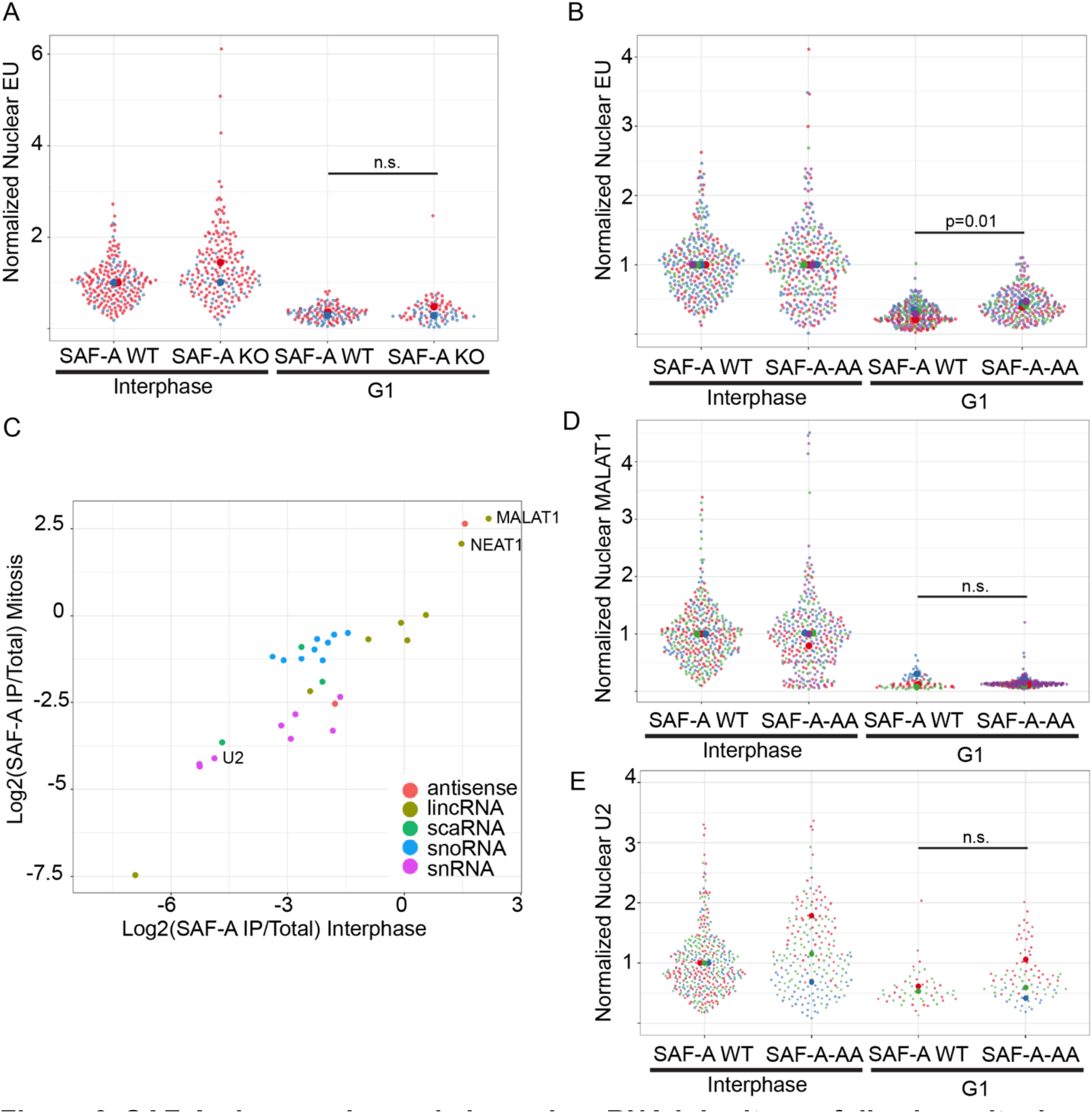
SAF-A plays a minor role in nuclear RNA inheritance following mitosis. A. SAF-A-AID WT (-IAA) and SAF-A-AID KO (+IAA) were synchronized in G2 and labeled with EU as described in Figure 1. Cells were then released from G2 and nuclear RNA was detected and quantified in G2 and G1 cells. SAF-A KO did not result in a significant change in nuclear RNA inheritance. B. SAF-A-AID KO (+IAA) cells expressing rescue constructs of SAF-WT or SAF-A-S14A-S26A were synchronized in G2 and labeled with EU as described in Figure 1. Cells were released into G1 and nuclear RNA labeling was quantified in G2 and G1 cells of each genotype. C. Conserved, chromatin-bound RNAs from Figure 2 were compared to enrichment in SAF-A IP-RNA-seq. Nuclear lncRNAs MALAT1 and Neat1 were highly enriched in SAFA IPs while other classes of nuclear RNA were underrepresented. D-E. SAF-A WT and SAF-A-AA cells were synchronized in G2 and released into G1 followed by detection of MALAT1 and U2 by RNA FISH. Quantitation of nuclear RNA FISH signal for both RNAs in G1 cells revealed that SAF-A-AA does not change the inheritance of either of these transcripts in G1. In all experiments a Wilcoxon-rank sum test was used to compare samples.

To understand which nuclear RNAs SAF-A could regulate during mitosis we merged the list of consistent chromatin-bound RNAs with our SAF-A RIP-seq data [6]. We identified two clear classes of transcripts. First, nuclear lncRNAs (NEAT1, MALAT1, and hnRNPU-as1) were enriched in SAF-A immunoprecipitations during both interphase and mitosis (Figure 3C). Second, snRNA, snoRNAs, and scaRNAs were all underrepresented in SAF-A immunoprecipitations (Figure 3C). This suggests that SAFA regulates the localization of nuclear lncRNAs but does not participate in regulation of other classes of nuclear RNAs.

To test the role of SAF-A in inheritance of different classes of nuclear RNAs we examined the G1 localization of U2 and MALAT1 in SAF-A-AA cells. Surprisingly, we detected no effect of expression of SAF-A-AA on the inheritance of either U2 or MALAT1 transcripts in G1 (Figure 3DE). Taken together, we conclude that SAF-A binds to nuclear lncRNAs that are not inherited after mitosis, and that depletion or forced retention of SAF-A on mitotic chromatin does not lead to an increase in the nuclear inheritance of specific transcripts.

### Dynamic SAF-A-AA binding to mitotic chromatin explains lack of effect on nuclear RNA inheritance

To understand why SAF-A-AA did not result in a more dramatic effect on nuclear inheritance of the MALAT1 transcript we examined the dynamics of SAF-A-AA-GFP during mitosis using FRAP. We degraded endogenous SAF-A-AID using auxin and simultaneously expressed SAF-A-AA-GFP or SAF-A-WT-GFP as the only source of SAF-A. Consistent with our previous work we found that ~40% of SAF-A-AA-GFP accumulated on the surface of mitotic chromosomes in living cells compared to ~8% of SAF-A-WT-GFP (Figure 4AB). To measure the dynamics of SAF-A-AA binding to mitotic chromosomes we performed FRAP of chromosomal regions in cells expressing SAF-A-AA-GFP and performed qualitative analysis of FRAP data [26]. Interestingly we found that SAF-A-AA rapidly recovered on the mitotic chromosome surface with a halftime of recovery of ~3 seconds and 96% of the total protein being part of the mobile fraction over the course of a 1 minute imaging experiment (Figure 4CD). We conclude that SAF-A-AA increases the residence time of SAF-A:RNA complexes on mitotic chromatin, but that SAF-A-AA is highly dynamic with the majority of the protein exchanging with the cytoplasmic pool. The highly dynamic exchange of SAF-A:RNA complexes between the chromosome and cytoplasm likely explains the lack of dramatic effects of SAF-A-AA on nuclear RNA inheritance of MALAT1 in G1, since an intact nuclear envelope in G1 would theoretically limit this mode of exchange.

**Figure 4.**
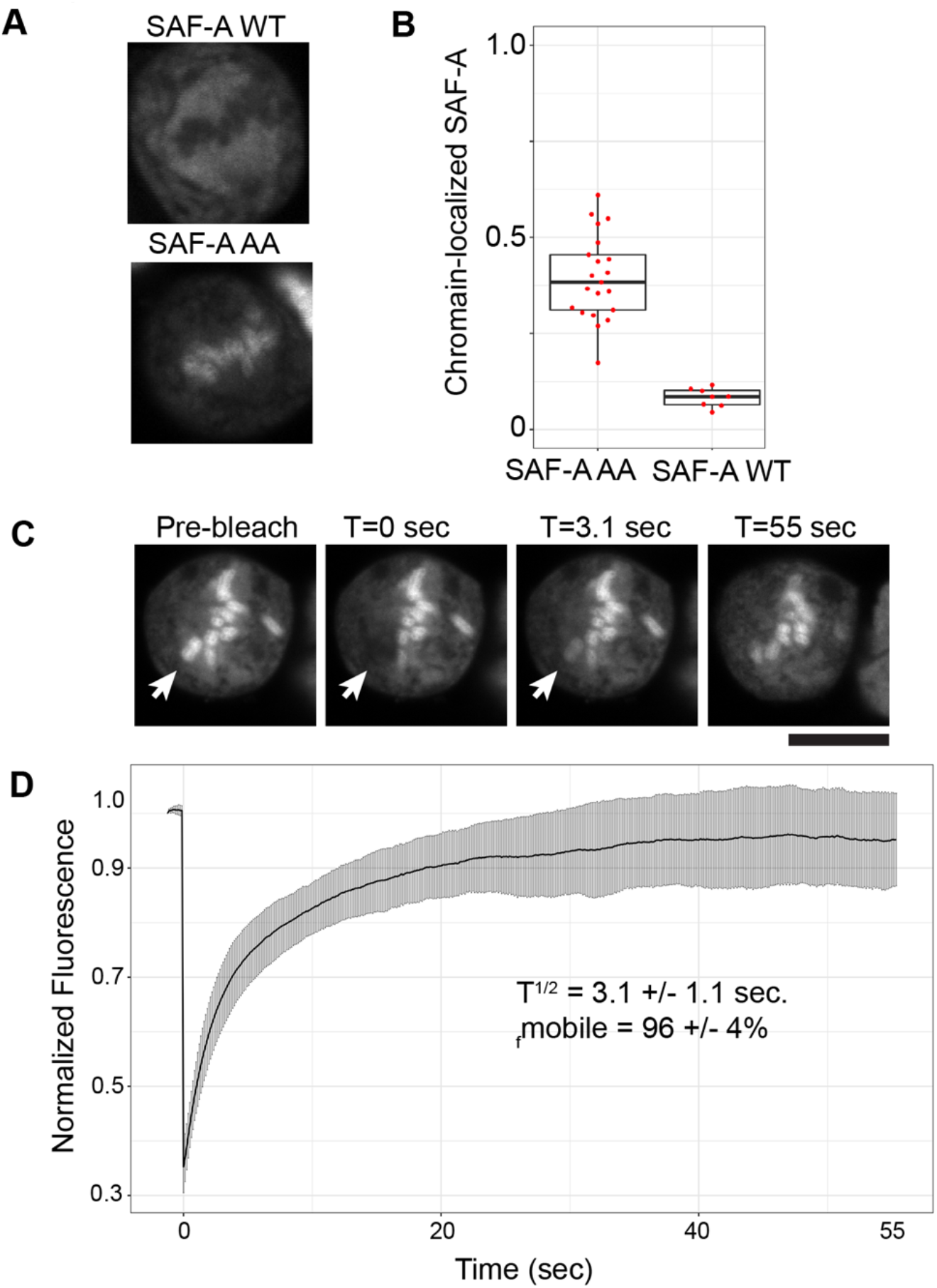
SAF-A-AA is highly dynamic on mitotic chromatin and enters the nucleus after nuclear envelope reformation. A. SAF-A-AID KO (+IAA7) expressing SAF-A-WT-GFP or SAF-A-AA-GFP were observed during mitosis using live cell imaging. SAF-A-AA outlined individual chromosomes while SAF-A-WT is excluded from chromosomes. B. Quantitation of the fraction of SAF-A WT or AA bound to chromosomes from live-cell imaging. C. The dynamics of SAF-A-AA-GFP were observed using FRAP over the course of ~ 1 minute. D. Average FRAP recovery of SAF-A-AA-GFP (n=25 cells) along with descriptive statistics of average FRAP behavior. Scale bars are 10 μm.

### SAF-A and U2 RNAs reenter the nucleus following nuclear envelope reformation

To understand whether nuclear RNA inheritance in G1 cells requires a nuclear reimport mechanism, we examined the chromatin localization of U2 RNA and SAF-A protein in early G1 cells relative to nuclear envelope reformation. We found that U2 RNA accumulates in G1 cells after nuclear envelope reformation (Figure 5A). We identified cells in very early G1 with partially reformed nuclear envelopes and cytoplasmic U2, but as soon as the nuclear envelope was sealed we observed complete localization of U2 to the nucleus. Interestingly, we identified a very similar pattern of localization with SAF-A. SAF-A was cytoplasmic in early G1 cells with partially reformed nuclear envelopes (Figure 5B), but complete nuclear localization of SAF-A in cells with an intact nuclear envelope. Taken together, these results demonstrate that nuclear RNA and RNA binding proteins accumulate in G1 cells following nuclear envelope reformation.

**Figure 5.**
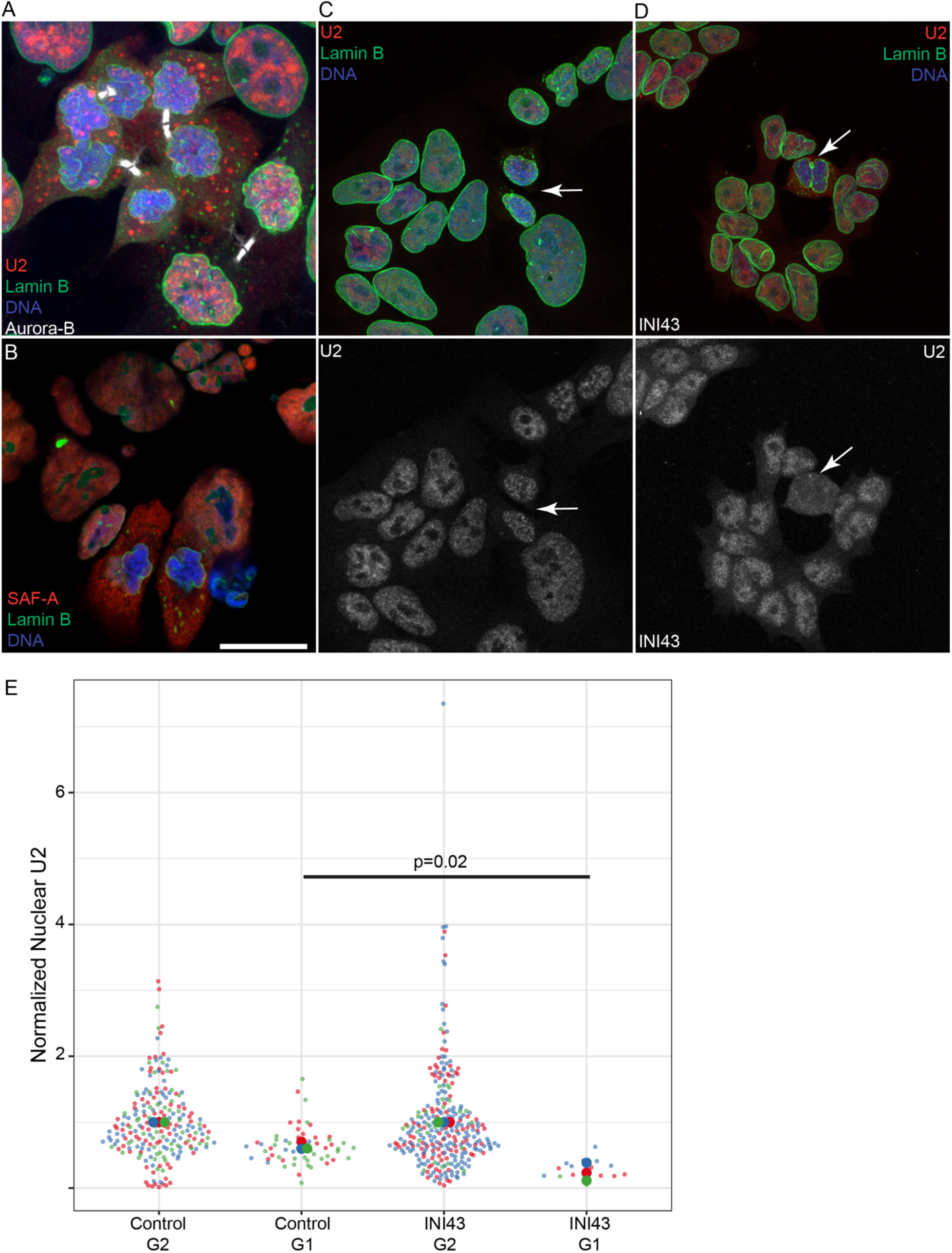
Nuclear import regulates RNA inheritance in early G1. A. U2 RNA FISH was conducted with staining for Aurora-B (white) and Lamin B. B. SAF-A was detected in early G1 cells with Lamin B. C-D. Cells were treated in mitosis with DMSO or INI43 and allowed to proceed into G1. U2 RNA, Lamin B, and Aurora-B were detected to identify early G1 cells (arrows). E. Quantification of nuclear U2 FISH signal in DMSO and INI43-treated cells. Different colored dots indicate biological replicates. Scale bars 10 μm.

### Nuclear import is required for U2 accumulation in G1 nuclei

To determine if nuclear import is required for the accumulation of U2 RNA in the nucleus in G1, we inhibited nuclear import using the small molecule INI43 [27]. Previous work has demonstrated that snRNPs are imported into the nucleus following maturation in an importin β-dependent manner [28]. INI43 was developed to directly target importin β and inhibits importin β-dependent transport [27]. We synchronized cells in G2 using RO3306, allowed them to enter into mitosis for 45 minutes, then added DMSO or INI43 to cells as they exited mitosis. We then examined to localization of U2 RNA by FISH relative to the nuclear envelope in early G1 cells. Consistent with our previous results we found that U2 RNA rapidly accumulated in early G1 cells treated with DMSO (Fig. 5C, E). In contrast, early G1 cells treated with INI43 exhibited a significant defect in U2 nuclear accumulation despite a closed nuclear envelope (Fig. 5DE). These results confirm our hypothesis that U2 RNA is inherited by daughter cells following mitosis through nuclear import after nuclear envelope reformation.

Prior to this study, RNA inheritance following mitosis has been queried for a limited number of specific transcripts. For example, previous work found that unprocessed rRNA is stored during mitosis and reimported into the nucleolus of G1 cells [15]. Unlike pre-rRNA, XIST RNA and repetitive RNAs do not appear to be inherited following mitosis [7, 8]. In this work, we devised a strategy to monitor nuclear RNA inheritance during late mitosis and found that a significant fraction of nuclear RNA is inherited by daughter cells in G1 following mitosis.

We found that a prominent example of an inherited RNA is the spliceosomal RNA U2. In contrast, long noncoding transcripts such as MALAT1 and NEAT1 are not inherited in G1 cells, but rather remain in the cytoplasm following cell division. Accumulation of MALAT1 and NEAT1 in the nucleus requires new transcription of these transcripts, which is reinitiated each cell cycle in G1. Interestingly, while mutation of SAF-A mitotic phosphorylation sites leads to a steady-state accumulation of RNA on mitotic chromosomes, SAF-A protein associates dynamically with mitotic chromatin leading to little effect on G1 RNA inheritance.

Taken together, our results support a model where distinct mechanisms regulate RNA dissociation from chromatin during mitosis and inheritance by G1 cells following division. In this model, SAF-A is responsible for removing the bulk of nuclear RNAs from chromatin during early mitosis. In late mitotic and early G1 stages, we speculate that small nuclear RNAs—such as snRNAs, snoRNAs, and scaRNAs—are inherited following division by reimportation of the RNPs into the nucleus. This interpretation is supported by the observation that each of these classes of RNA have extremely long half-lives [29]. In contrast, we observe that lncRNAs that associate with chromatin through SAF-A are removed from chromatin during mitosis and are not inherited in G1. Because SAF-A does reenter the nucleus in late mitosis, these transcripts must be stripped from SAF-A and remain in the G1 cytoplasm. It is possible that differential nuclear import of transcripts following mitosis is a result of size differences in nuclear RNA or the fact that lncRNAs do not normally exit the nucleus and do not have an established nuclear import pathway. It is tempting to speculate that mitosis serves as a mechanism to remove lncRNAs from chromatin and increase transcript degradation rates. It is possible that removal of lncRNAs from chromatin allows cells to change transcriptional states following mitosis.

---

**Supplemental Figure 1.**
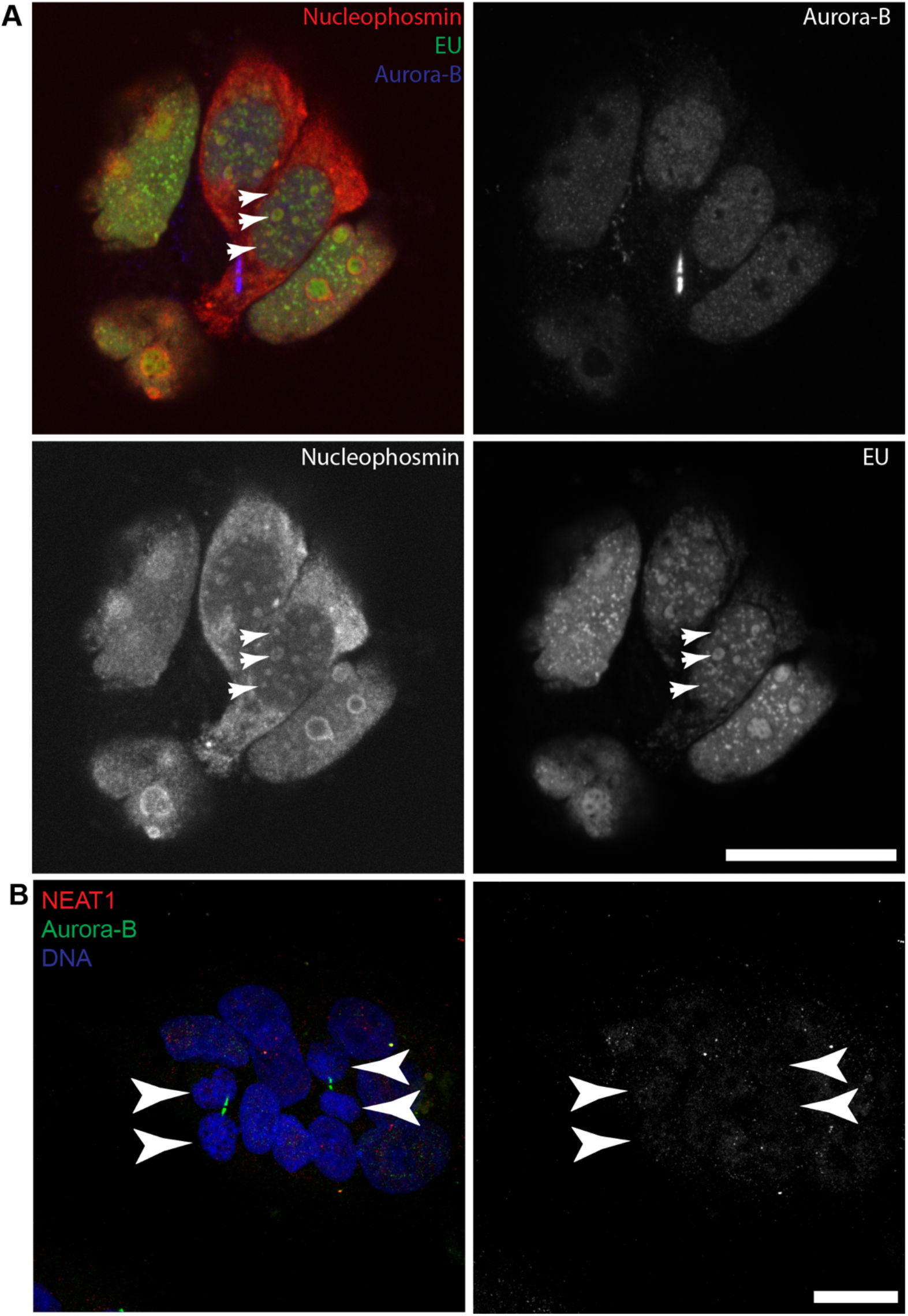
Nucleolar RNA is inherited in G1. **A.** Codetection of EU (labeled in G2 cells) and Nucleophosmin in G1 cells. EU RNA forms irregular chunks in G1 nuclei (arrowheads) that partially colocalize with the nucleolar marker Nucleophosmin. B. RNA FISH detection of NEAT1 RNA in interphase and early G1 cells (arrowheads). Scale bars ar 5 μm.

**Supplemental Figure 2.**
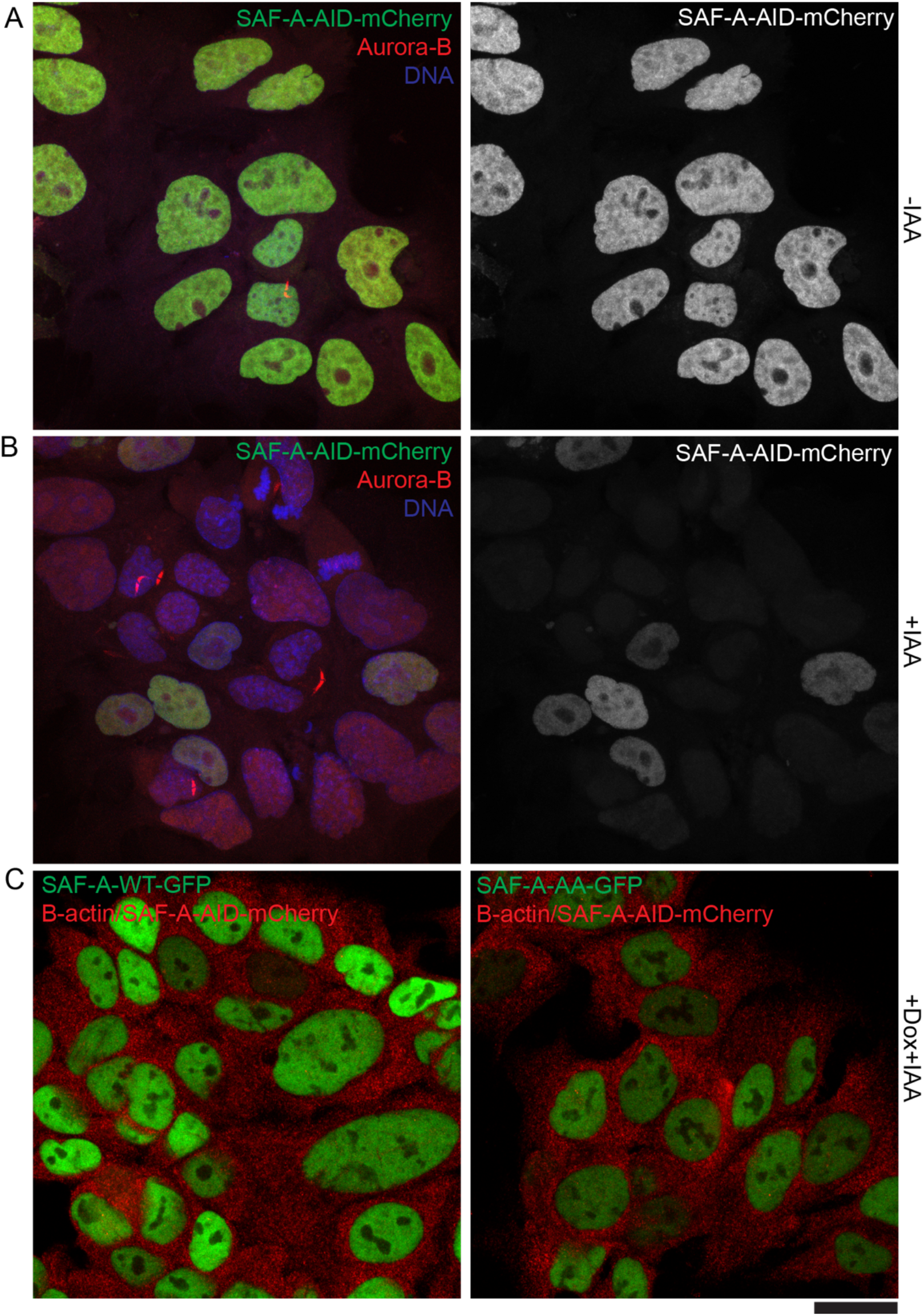
Validation of SAF-A cell lines. A-B. Fluorescence of SAF-A-AID-mCherry in SAF-A-AID cells with and without IAA addition. C. Fluorescence of SAF-A-GFP rescue transgenes in SAF-A-AID-mCherry cells depleted of endogenous SAF-A and rescued with the indicated constructs. Endogenous SAF-A-AID-mCherry would appear in the red channel and B-actin FISH outlines the cytoplasm. Scale bar is 5 μm.

## Materials and Methods

### Cell culture

DLD-1 cells and DLD-1 cell derivatives were cultured in RPMI-1640 (Sigma) + 10% FBS (Cytivia) + Pen/Strep. DLD-1 derivatives used in this study were: DLD-1 (SAF-A mAIDmCherry Bsr + OsTir1 Puro), (SAF-A-mAID-mCherry Bsr + OsTir1 Puro + Tet-ON-SAF-A-WT-GFP Puro), (SAF-A-mAID-mCherry Bsr + OsTir1 Puro + Tet-On SAF-A-S14A-S26A-GFP Puro). To induce SAF-A degradation cells were treated with 1 μM doxycycline and 1 μM IAA for 2 hours. We found that longer SAF-A depletion times resulted in dramatic reduction of nuclear transcripts as detected by 5EU labeling (not shown). To induce SAF-A allele exchange, cells were incubated cells with 1 μM Doxycycline and 1 μM IAA for 24 hours. Cell synchronization was accomplished by treatment 8 μM RO3306 for 24 hours. Transcription inhibition was accomplished by arresting cells in RO3306 for 24 hours, followed by RO washout for 30 minutes, followed by the addition of either 1 μM triptolide, 1 μM THZ1, 1 μM LDC, or 15 μM INI43 for 1.5 or 2 hours. Cells were incubated with 1 mM 5EU as described in each figure.

### Cell fixation and combined immunofluorescence and FISH

Following cell synchronization cells were fixed with 4% PFA (Electron Microscopy Services) in PBS for 10 minutes at room temperature. Cells were then permeabilized by incubation in PBS + 0.5% Triton-X100 from 20 minutes at room temperature. Cells were then stored in 70% ethanol until use. For immunofluorescence the following antibodies were used: mouse anti-Aurora-B (BD Bioscience), mouse anti-lamin B (Abcam: Ab16048). Cells were rehydrated in PBS + 0.1% Tx-100 for 30 minutes, incubated in PBS-T + 5% ultra pure BSA + RNasin for 30 minutes. Cells were incubated by primary antibody for 1h at 37C or 3h at room temperature. Cells were washed 3X 5 minutes with PBS-T, then incubated with secondary antibody for 1 hour. Cells were then washed 3X 5 minutes with PBS-T. Cells were postfixed with 2% PFA in PBS for 10 minutes at room temperature. Cells were then blocked in 2XSSC 50% formamide + 5% ultra pure BSA + RNasin for 30 minutes to 2 hours at 37C. Cells were incubated with FISH probes overnight and washed essentially as described [31]. EU was detected essentially as described [6].

### Fluorescence recovery after photobleaching

Cells expressing either SAF-A-WT-GFP or SAF-A-S14A-S26A-GFP were seeded onto glass bottom 35mm dishes and allowed to grow for 24 hours. Cells were imaged on a Nikon A1R confocal microscope equipped with a stage top incubator and CO2 chamber (Tokai Hit). Mitotic cells were selected through the oculars and selected for FRAP. FRAP was performed with 50% laser power using the 405 laser for 1 second in a 4 μM x 4 μM square. Images post recovery were acquired every 0.06 seconds at 256 x 256 pixel resolution using a scan zoom setting of 8. Post recovery images were acquired for 45 seconds after bleaching. A total of 20 mitotic cells expressing SAF-A-AA were acquired and 10 mitotic cells expressing SAF-A-WT. Bleaching correction and qualitative analysis of FRAP recovery kinetics were performed as described [26].

### Image acquisition and analysis

All images were acquired using a Nikon A1R confocal microscope equipped with a 60X 1.4NA Vc lens and laser lines at 405, 488, 562 and 647nm. Images were collected using a scan zoom of 2X (140nm/pixel) using a galvano scanner. Images were acquired a Z stacks spaced 0.3 μM apart. Each experiment was repeated in at least two biological replicates (precise N or replicates in indicated in figure legends). Between 5-10 fields of cells (~100-200 cells) were acquired for each condition. Fluorescence intensity of EU or RNA FISH signal was measured in the nucleus of each cells using a script in ImageJ. Cells in telophase or G1 were manually identified an annotated using Aurora-B staining. For each experiment fluorescence intensity was normalized to the average intensity of untreated G2-arrested cells. Statistical comparison of conditions was performed using a Wilcoxon Rank Sum test. SuperPlots of normalized measurements for each experiment were produced using R [30].

### Analysis of chromatin-bound RNAs and SAF-A enrichment

Published lists of chromatin-bound RNAs were downloaded from published articles. Intersection of the chromatin-bound RNAs were calculated by merging lists using gene names. Genes that appeared in three conditions were selected for further analysis. Chromatin bound RNAs were intersected with SAF-A enriched RNAs using gene names. Plots of chromatin RNA analysis were prepared using R.

## Acknowledgements

This work was supported by the NIH (GM122893-06 to M.D.B.). The authors thank members of the Blower lab for helpful discussions and comments on the manuscript.

